# Bamboos flower after the return of almost the same sun-moon phasing as at seedling emergence

**DOI:** 10.1101/2021.06.11.448081

**Authors:** Benoit Clerget

**Affiliations:** CIRAD, UMR AGAP Institut, Univ Montpellier, 34398 Montpellier, France

**Keywords:** bamboo flowering, *Bambusoideae*, flowering time, lunar phases, photoperiod, solar year

## Abstract

All Asian woody bamboo species of economic importance are semelparous. They remain vegetative during time intervals that are specific to each species and range from three to 120 years, with notable concentrations around a series of values (3, 7-8, 14-17, 29-36, 42-48, 61-64, and 120 years). Then, they flower gregariously within a short period. Entire forests temporarily disappear during these periods, and the dates of these dramatic events have been recorded over the last 200 years.

While searching for a correlated environmental cue, I have found that the concentrations of flowering cycles were highly correlated with the series of successive returns of almost the same sun-moon phasing as at seedling emergence. On basis of knowledge on plant photoperiod sensitivity, I hypothesize that bamboo plants i) run a lunar cellular clock that is set at the full moon, ii) retain in their cellular memory the exact sun-moon phasing of the year of their emergence as seedlings, and iii) inhibit flowering until the occurrence of a unique, species-specific sun-moon phasing that is shifted by a precise amount from the sun-moon phasing at their emergence. Recent evidence of plant responses to lunar cycles supports this hypothesis, for which experimental evidence is now anticipated.

**Highlight:** The long-standing enigma of the triggering of the gregarious flowering of bamboos is possibly solved. Flowering would be inhibited until the occurrence of the species-specific sun-moon phasing.

## Introduction

Bamboos constitute a group of 1642 species belonging to the grass family (Poaceae) that are distributed on four continents, excluding Europe (Zheng *et al*., 2020). They exhibit either a ‘clumping’ growth form, with short rhizomes that simply expand the root network, or a ‘running’ growth form, with long rhizomes that can aggressively expand underground. The culm stock is permanently renewed by the annual emergence of new culms from the rhizomes during a specific time window and the death of old culms (Zhao-hua *et al*., 2005). The natural turnover period of culms is 7-8 years. Culms reach their maximum height and width within a few weeks; thus, in species that achieve heights of 20 to 30 m, their daily height growth exceeds 1 m. One group of bamboo species flower continually or yearly, and other species flower sporadically, i.e., after irregular intervals of several years. The latter group of species is semelparous, with periodic gregarious monocarpic flowering; entire monospecific forests of such bamboos flower simultaneously after a set number of years, undergo mast-seeding, die, and regenerate from new seedlings. All Asian bamboo species of major economic importance are mast-seeding species (Janzen, 1976). Bamboo is used as food, material for construction or furniture, fuel, fibre for textiles and paper production, and kitchenware (Troya Mera and Xu, 2014; Takano *et al*., 2017). Wild animals, such as the giant panda, are also affected by bamboo cycles (Janzen, 1976; Campbell and Qin, 1983; Zheng *et al*., 2020).

One hundred Asian bamboo species have been used and cultivated for centuries. Because of the importance of these mast-seeding events, historical records of the years in which they occurred have been kept since at least 833 and mainly since the middle of the nineteenth century. A preliminary data collection effort provided usable information from old records about the mass flowering years and the flowering cycles (the durations between two consecutive flowering years) for 33 species from South Asia (Janzen, 1976). A notable concentration of mean periods about multiples of 15-16 years (3, 7-8, 14-17, 29-36, 42-48, 61-64, and 120 years) was then noticed within these data as well as data for four Chinese bamboo species (Campbell, 1985). This series of multiple values could have resulted from two evolutionary series derived from two ancestors with flowering cycles of 2 and 5 years (Veller *et al*., 2015). This list was recently increased to 85 species by adding observations from China, South America and Australia (Zheng *et al*., 2020) (Supplementary Table S1 at JXB online).

Correlations between the bamboo flowering cycles and environmental conditions were sought in order to determine the factor responsible for these concentrations of long to very long flowering cycles. Weak relationships were established with (1) the 11-year sunspot cycle for North American Arundinaria gigantea species (Campbell, 1985), (2) a 30-40 year cycle of strong drought years in East Asia with a general phase shift from North to South, (3) a 45-year drought cycle in India, (4) a 7-year drought cycle in the Southern Hemisphere and in South India, (5) wet years in SW India, Jamaica and SE USA12, and (6) a 100-year cycle of strong earthquakes in China (Campbell and Qin, 1983). However, all of these correlations lack convincing generality (Franklin, 2004).

I tested the hypothesis of a possible relationship between bamboo flowering cycles and lunar cycles (29.53 days in average) and found the very high correlation presented in this paper.

## Material and methods

The recently published updated list of the mean flowering cycle of 85 bamboo species was used in this study (Zheng *et al*., 2020) (Supplementary Table S1). Data for the species *Phyllostachys bambusoides* from a previous list were added (Janzen, 1976). When the range of values for a cycle was lower than ten years, the mean value of the cycle was used. Otherwise, two or more values were taken for the species when the range was higher than ten years, thus indicating variability in the flowering cycle within the species (Alam, 2008). Thus, 108 mean flowering cycles observed in 86 species were analysed.

The shift in the sun and moon phases after exactly one year was computed using a year duration of 365.2563631 days (International Earth Rotation and Reference System Service, 2010) and a mean synodic lunar cycle of 29.53058886 days (Seidelmann, 1992). The computation started at day_i_ of year 0 and was yearly run. A list of 19 years over a period of 160 years when the moon phase at day_i_ differed by less than two days from the original phase was established, and two additional values with differences less than 3.2 days, at 8 and 16 years, were added.

Automatic clustering of the 108 mean observations was performed using the MODECLUS procedure (k=4) in SAS (SAS, 2012). A second supervised clustering was performed in Excel by aggregating each observed flowering cycle to the closest value among the values on the list of 21 years with appropriate moon phasing.

Only basic statistics can be done on such a dataset, in which sampling is not expected to follow a specific distribution. The cluster means were regressed against the observed values with the SAS procedure REG to calculate the R^2^ with the model probability. The root mean square difference (RMSD) was calculated, and a paired, two-sided t-test was performed to compare the two means using the SAS procedure TTEST on the same set of values. Finally, the same statistics were applied to the observed values compared against the year values from the list of appropriate years used in the supervised clustering process.

## Results

Starting from any solar day (day_i_) on year 0, the lunar phase (in the number of days since the new moon) at dayiof the following years irregularly oscillates in 2-3 year cycles (Fig. 1). In some years, the lunar phase at day_i_ is close to either 0 or 29.53 days, i.e., it varies from the original phase by less than two days. After 160 years, the moon phase difference at day_i_ reaches a minimum value of −24 min, and a new series starts again for the next 160 years, with very similar values to those of the previous cycle (Fig. 2). Negative values indicate an advance of the moon phase at day_i_ when compared with year 0. Among the shift values of less than two days, this advance regularly decreased from −1.5 days to −24 min in a (8-27-46-65-84-103-122-141-160) series of years; this series had notable similarities with the previously described concentrations of flowering cycles (3, 7-8, 14-17, 29-36, 42-48, 61-64, 120 years) (Campbell, 1985; Veller *et al*., 2015). Similarly, positive values of the phase shift regularly increased from 4 h 22 min to 1.8 days in a (19-38-57-76-95-114-133-152-11-30) series of years that completed the series with negative day-shift values.

**Fig. 1.**
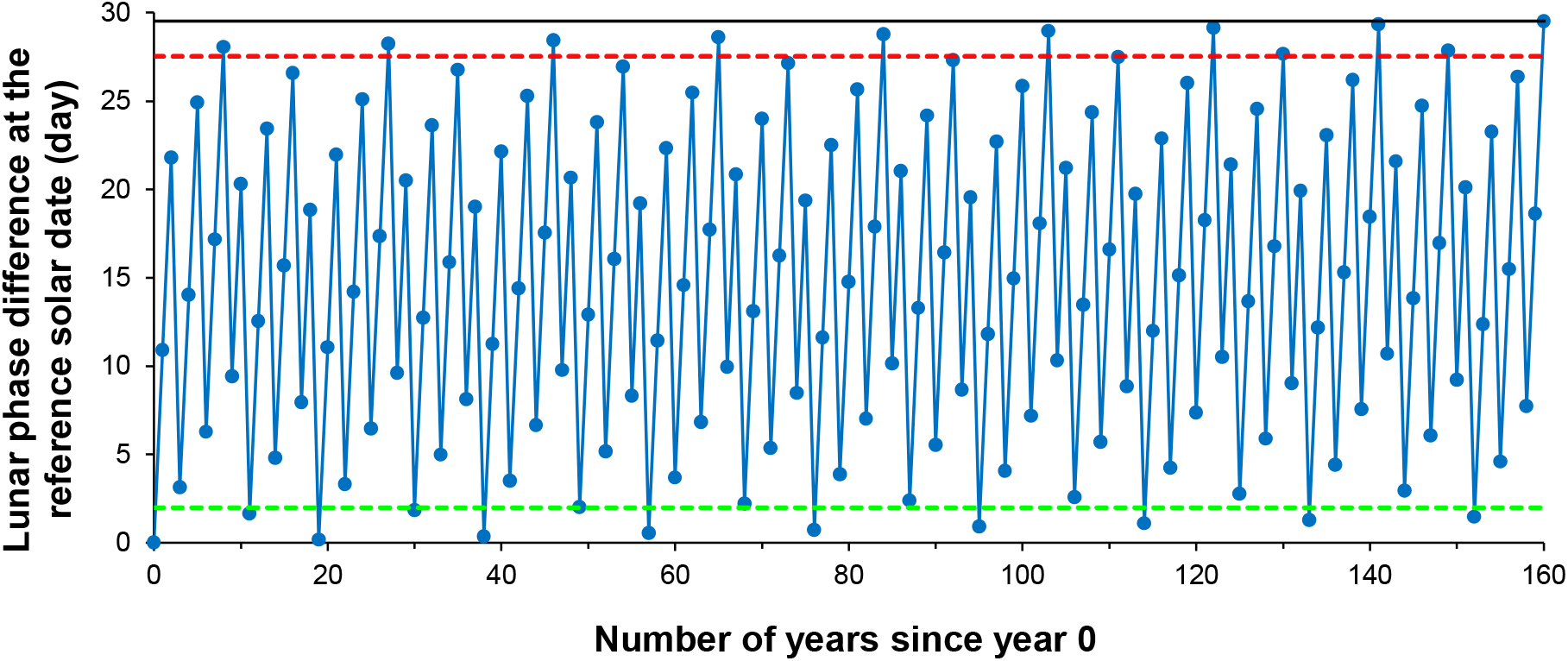
Yearly moon phase shift from the initial situation. The moon phase at exactly the same solar position oscillates with years. Red and green dot lines indicate lower than two days-differences with the situation on year 0.

**Fig. 2.**
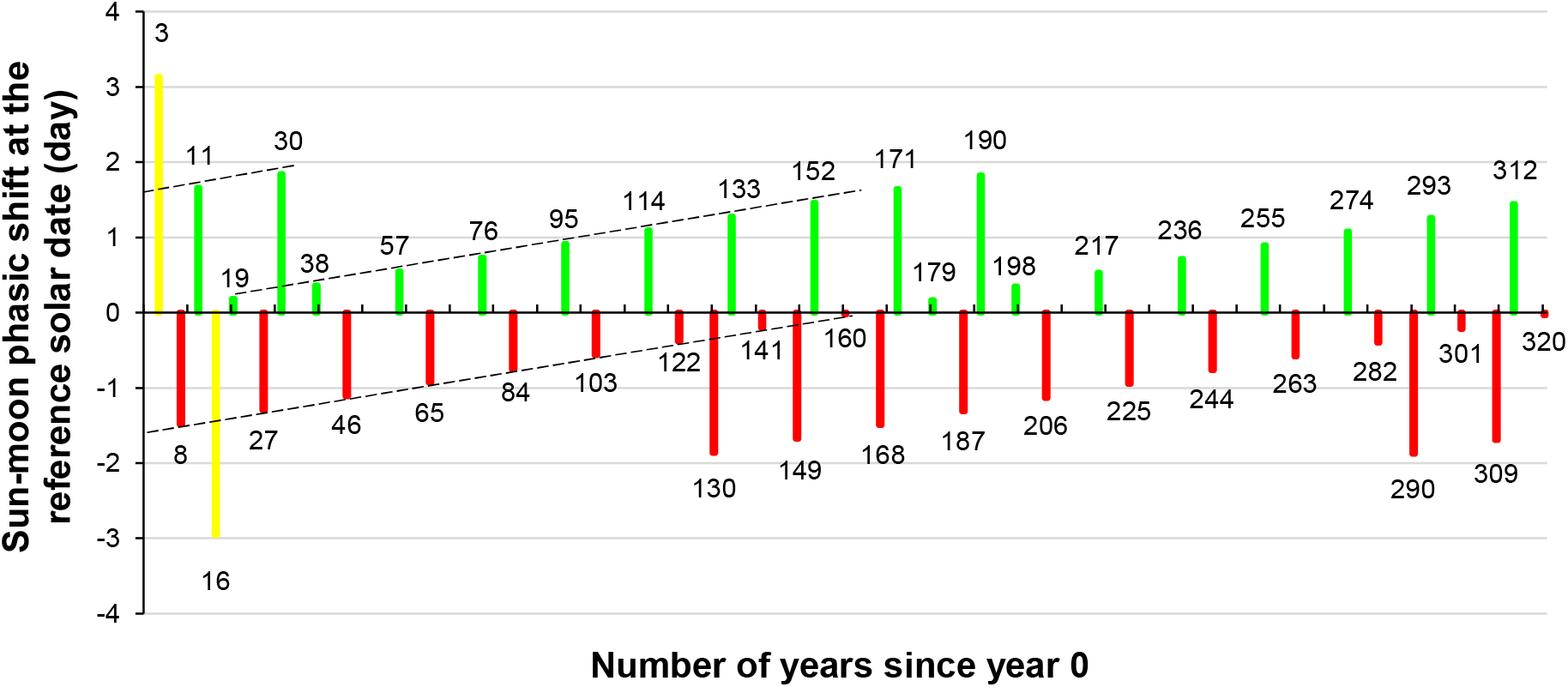
Years with sun-moon phasing differing by less than two days from the original situation. Moon phase is ahead of the original situation in the green series and lagging behind it in the red series. Two values with a threshold value of 3.2 days difference are added in yellow.

**Fig. 3.**
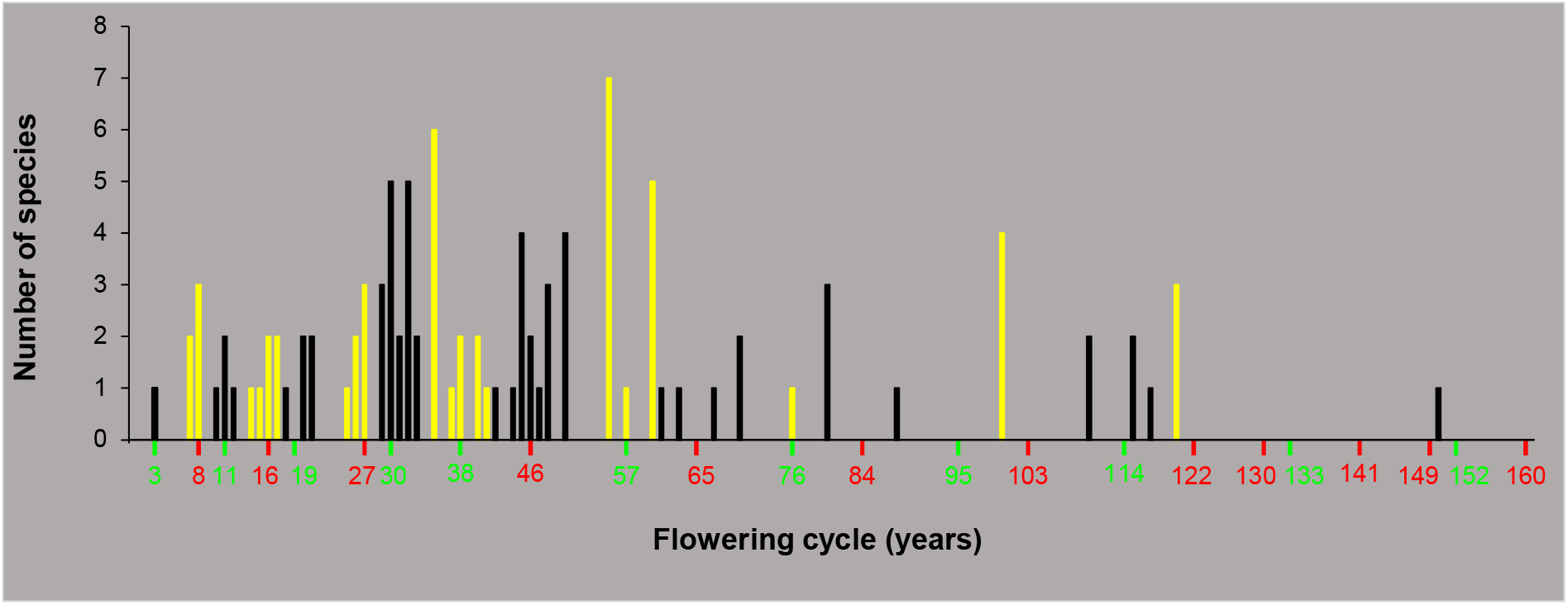
Distribution of the 108 flowering cycles grouped in 17 clusters. Clusters are alternately in black and yellow, after aggregating the observations by closest distance to one value in the list of years with appropriate sun-moon phasing. The years of the return of an appropriate sunmoon phasing are tagged either in red or in green.

Values close to the observed values of 3 or 14-17 were missing from these two series. However, in years 3 and 16, the phase difference was lower than 3.2 days, and these two values were consequently added to the series of years when the moon synchronization was apparently correlated with the bamboo flowering cycles. In the final list, 21 years during the 160 years had an appropriate moon phase at day_i_ (3-8-11-16-19-27-30-38-46-57-65-76-84-95-103-114-122-133-141-152-160), and these 21 years occurred 7.6 years apart, on average.

The automated clustering of the mean durations of the 108 observed flowering cycles resulted in a series of 17 clusters centred on (3-8-11-16-21-26-30-32-36-41-46-50-55-63-81-100-116) (R^2^=0.996 with P=0; RMSD=2.01; t-test =0.52 with P=0.60) (Supplementary Fig. S1). Until 16 years, the clusters perfectly fit the concentrations of flowering cycles. However, from 21 to 50 years, the automatic method produced height clusters rather than the two reported concentrations. The observation density was higher within this range of durations than elsewhere, and the automatic process split possibly homogenous clusters, as observed around year 30 (Supplementary Fig. S1).

For these reasons, supervised clustering was then performed by grouping the data based on the shortest distance to the 21 values from the list of years with appropriate sun-moon phasing; this approach also resulted in 17 clusters (Fig 1). The statistics were close to those of the automated clustering, with a more significant t-test result (R^2^=0.996 with P=0; RMSD=2.01; t-test =0 with P=1). P values were very high when using the mean flowering cycles and the 107 degrees of liberty among means. Thus, individual observations, which are not always accessible, would not improve the result. When the years from the list of right phasing events instead of the cluster means were used, the statistics were slightly less accurate due to the distance between the two values (R^2^=0.995 with P=0; RMSD=2.24; t-test =0.42 with P=0.67).

A 122-year flowering cycle was applied to the flowering of *Phyllostachys bambusoides* recorded in 1114 (Janzen, 1976) and the flowering of *Phyllostachys nigra* var. *henonis* recorded in 813 (Table 1). The predicted flowering years fit a large proportion of the observations well. On the other hand, semelparity is found in a wide range of plant species covering at least 20 families (Sharma *et al*., 2008). *Strobilanthes kunthianus*, a shrub from southwestern India, is among these species, and it has rigorously flowered once every 12 years since the first recorded flowering event in 1838. Flowering in this species occurs in September-October, and seed set occurs at the end of the year, after the dry season begins. Seeds germinate at the onset of the monsoon rains in May of the following year. Thus, they flower 11 years later and complete a 12-year flowering cycle, which perfectly fits with one of the values on the list of years with appropriate moon phasing.

**Table 1.** Flowering years of two bamboo species. Data recorded in Japan since 999 and 813 and predicted flowering years based on a flowering cycle of 122 years applied from 1114 and 813.

| *Phyllostachis bambusoides* |  |  | *Phyllostachis nigra* var. *henonis* |  |  |
| --- | --- | --- | --- | --- | --- |
| Recorded | Predicted | Other cycle | Recorded | Predicted | Other cycle |
| 999 |  | 115 | 813 | 813 |  |
| 1114 | 1114 |  | 931 | 935 |  |
|  | 1236 |  |  | 1057 |  |
|  | 1358 |  |  | 1179 |  |
|  | 1480 |  | 1247 |  | 68 |
|  | 1602 |  |  | 1301 |  |
| 1715-1736 | 1724 |  |  | 1423 |  |
| 1844-1847 | 1846 |  |  | 1545 |  |
| late 1960's | 1968 |  | 1666 | 1667 |  |
|  |  |  | 1786 | 1789 |  |
|  |  |  | 1848 |  | 62 |
|  |  |  | 1908 | 1911 |  |

## Discussion

Each bamboo species and *Strobilanthes kunthianus* flower on dates that are close to the year of one of the returns of a sun-moon phasing similar to the phasing at seedling emergence. The correlation between the flowering cycles and the sun-moon appropriate phase calendar is very high, and a causal relationship seems very probable. Experimental evidence is now anticipated.

### How reasonable is a lunar hypothesis?

The lunar influence on animal, human, and plant activity has been a controversial topic for a very long time (Mayoral and Solbes, 2020). Because of tidal movements, the lunar influence on organisms living in the tidal zone is fully recognized, as in the case of the emergence of palolo worms on the Samoan Islands (Caspers, 1984). In contrast, many popular beliefs about lunar effects have not been statistically confirmed when studied, leading to scepticism in the scientific community. However, evidence confirming that lunar phases are perceived by animals and plants, which use them to synchronize their activity, is mounting. For example, a new chamber wall is deposited after a regular number of ~30 daily growth lines on the shells of nautiluses (Kahn and Pompea, 1978). Nautiluses are living fossils; they are benthic animals that have been living between −200 and −700 m depth in tropical oceans for 500 million years. In Nautiloid fossils, the number of growth lines increased as the number of lunar months increased during geological times. Thus, nautiluses perceive the lunar month in the dark depths of the oceans. Similarly, zooplankton follow rhythmic migration from the surface to − 50 m depth in phase with lunar day and month durations during Arctic winter, when the night is continuous (Last *et al*., 2016). The zooplankton would perceive the moonlight and respond to its changing intensity. However, moon tides generate internal waves of various types and frequencies inside fluid oceans that can be perceived by living organisms (Vic *et al*., 2019).

In contrast, atmospheric movements result mainly from chaotic meteorological effects. Thus, lighting variations are the only certain environmental signal of the passage of the time and the seasons on land (Thomas, 1998). In plants, it was recently shown that the daily rate of growth of sphagnum mosses on the northern Russian tundra exhibits a circalunar rhythm and is reduced by 30 to 50% at the full moon (Mironov and Kondratev, 2020). Moreover, Breitler *et al*. (2020) found an unusual midnight peak of expression of the key core clock gene LHY in coffee trees exposed to full moon light in a greenhouse. Further experiments showed that nightly artificial lighting at full moon luminance affected the expression of 3387 genes related to the circadian clock and the circadian regulation of the metabolism of coffee trees. Thus, minimal but significant evidence supports the hypothesis of the perception of the lunar cycle by bamboos and of its role in the control of their flowering cycle.

### Hypothesis of a circalunar internal clock

The possibility of a response to specific external years with a particular moon phasing on a specific solar date (the equinox, for instance) was examined and discarded. Indeed, after the first return of a specific sun-moon phasing needed for a specific species, no more similar events will occur until the end of the 160-year cycle. A biological reset after each flowering is thus required.

Thus, I hypothesize that bamboos run a lunar clock in addition to the previously identified clocks; the lunar clock is probably set by the full moon light due to the weak moon luminance (Breitler *et al*., 2020; Mironov and Kondratev, 2020). As suggested by Fisahn *et al*., (2012), a clock with a 24.8 h cycle (one lunar synodic day), set by the moon light instead of the tidal movement of Earth’s crust, would return to the same phase adjustment with the circadian clock after each lunar month. Based on the other clocks, bamboos recognize the times of the equinoxes (similar day-night duration) and of the solstices (stable sunrise and sunset times) (Borchert *et al*., 2005; Shim *et al*., 2016; Madrid *et al*., 2021; Clerget *et al*., 2021).These are four yearly reference dates with which plants can compare the sun and moon phases. In addition, bamboos must retain the exact memory of the lunar phase on a reference date during their year of the beginning of their life. This time could be either during seed formation or, more likely, at seedling emergence, which is known to be the time when plant developmental parameters are established (Kirby *et al*., 1985; Birch *et al*., 1998). In the case of *Strobilanthes kunthianus*, the lunar clock must be set at seedling emergence in order for the hypothesis to fit with the observations (Sharma *et al*., 2008).

In semelparous species, flowering is inhibited until the return of a sun-moon phasic correspondence that is sufficiently close to that at seedling emergence, but the requirement is even more precise. In each species, the long-term flowering inhibition ends after the occurrence of a specific sun-moon phasic adjustment, when the moon phase is either ahead of or lagging behind the original situation by a precise number of hours. The precision is less than four to five hours, which is the phase difference between two successive points in the series of appropriate years (Fig. 2). Lunar inhibition removal can occur one year before the flowering time and cause the inhibition of new culm outgrowth (Janzen, 1976; Alam, 2008). Although related evidence has not been systematically reported, bamboos are photoperiod-sensitive. In each observed species, new culms emerge during a specific period of the year, and flowering also starts in a specific month (Supplementary Table S1). Subtropical and temperate species from China, Japan and the Himalayan areas are long-day species that flower during spring between January and June. In contrast, tropical species from South and Southeast Asia are short-day species that flower between July and December. Additionally, orthologous genes of the highly conserved genes related to the circadian clock and floral pathways were identified in bamboos (Biswas *et al*., 2016; Xiao *et al*., 2018). Photoperiod-sensitive plants use thresholds in the daily solar period (daylength and daily changes in sunrise and sunset times) to temporarily inhibit the flowering process until a variety-specific solar day has been reached (Major, 1980; Thomas and Vince-Prue, 1997; Carberry *et al*., 2001; Clerget *et al*., 2021). Thus, after the removal of lunar inhibition, flowering is still inhibited by the photoperiodic response until the specific date in the year for the initiation of flowering in the species. The variability of these two systems controlling the flowering date within each species is unknown.

Experimental evidence is now needed to confirm that semelparous bamboos flower in response to a year with appropriate moon phasing. The long duration of the flowering cycles and the height of bamboo plants complicate this task. A gene network that oscillates with either a 24.8-hour or a 29.5-day period may be identified soon.

### Evolutionary considerations

Semelparous species probably evolved from annual photoperiod-sensitive monocarpic species that developed a way to use an additional lighting cue to further inhibit their flowering time through the already-existing gene network. They consequently remained monocarpic but gained substantial longevity, which allowed them to grow as tall as trees. This evolutionary innovation was successful and allowed bamboo to conquer vast areas in sufficiently humid zones from the equator to temperate latitudes. The possible selective advantages of mast seeding have been widely discussed. There are two main hypotheses: 1) predator satiety, which proposes that irregular fruiting cycles maintain the populations of seed predators at levels that prevent them from decimating a “mast year” of fruiting (Janzen, 1976), and 2) parental competition, which postulates that adult bamboo plants die to remove the intense shade that they often cast so that seedlings can become established (Janzen, 1976). The fire cycle hypothesis, which postulates the co-destruction of less fire-resistant emerging tree seedlings, is an additional possibility (Keeley and Bond, 1999).

## Supplementary material

Supplementary material were added at the end of the paper.

## Supplementary material

**Supplementary fig. S1.**
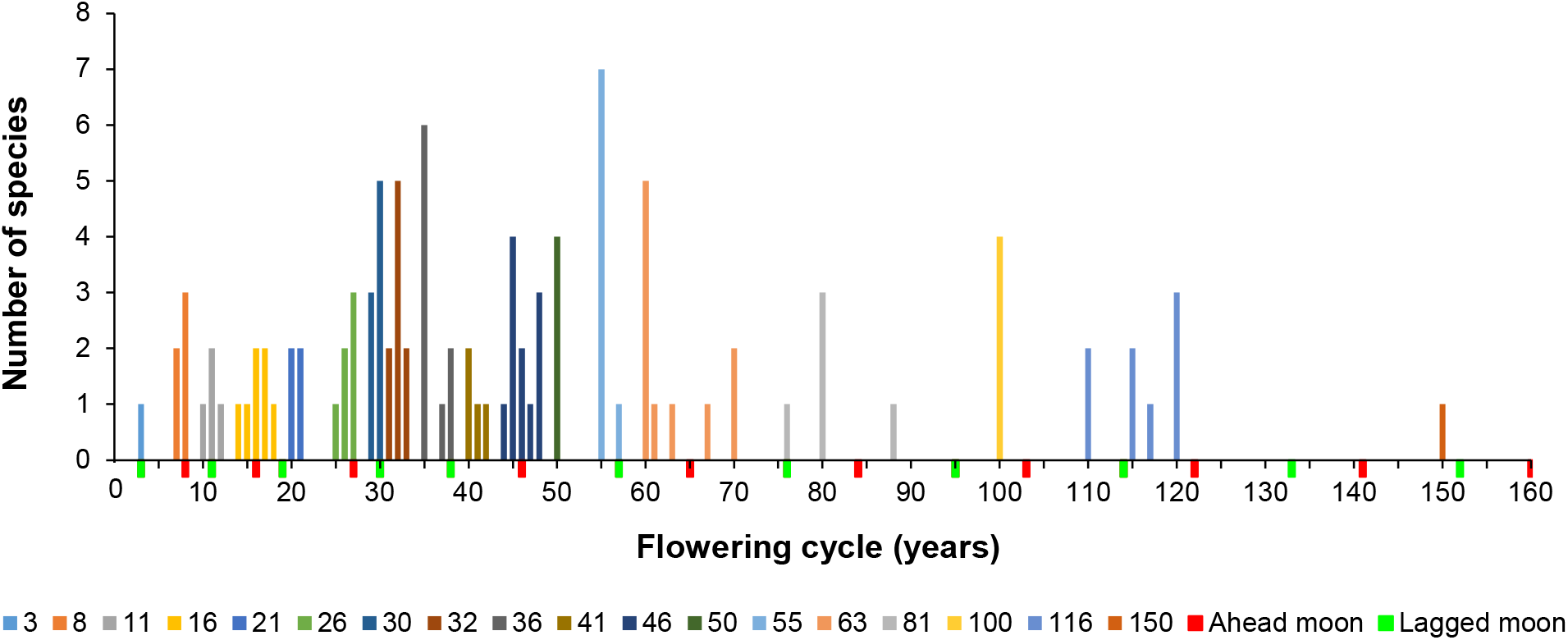
Distribution of the 108 flowering cycles automatically grouped in 18 clusters. The years of the return of nearly the same sun-moon phasing as at birth are tagged either in red (moon is a little ahead) or in green (moon is a little lagged).

**Supplementary table S1.**
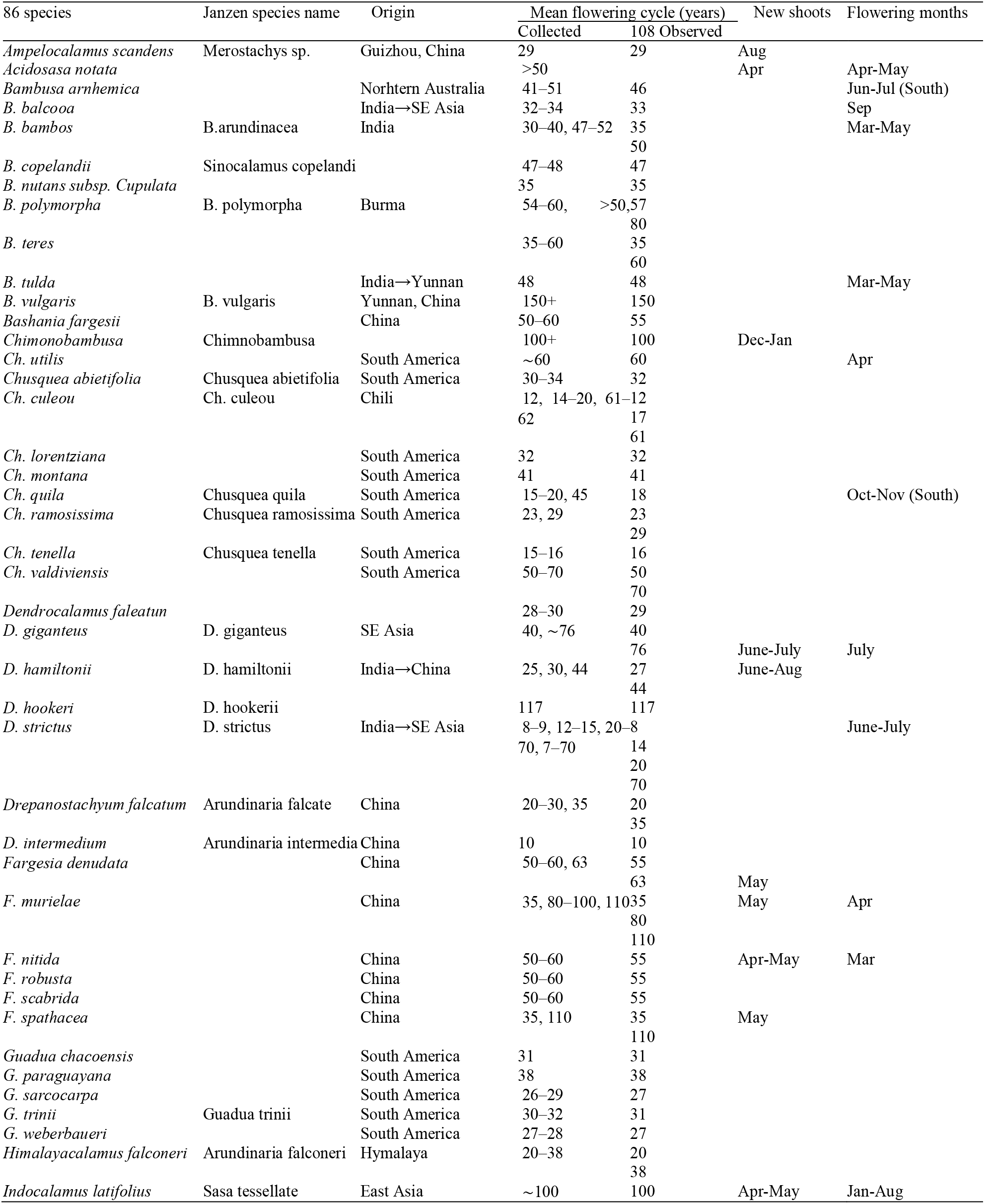
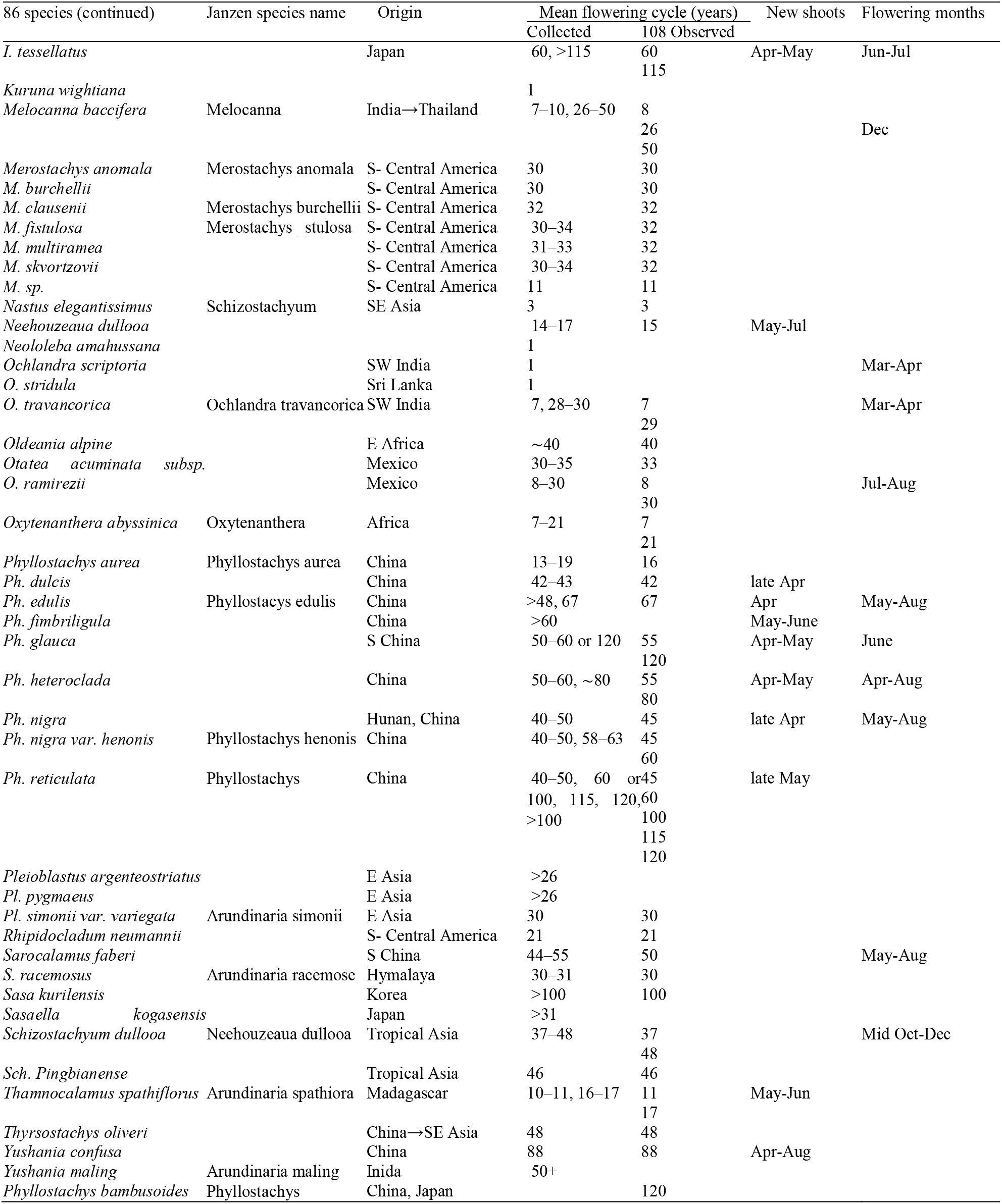
List of the 86 bamboo species. Origin, mean duration of flowering cycle, months of emergence of the new shoots and months of start and peaking of flowering are reported.

## Notes

### Competing Interest Statement

The authors have declared no competing interest.

### Summary of Updates

The title is more precised. Keywords were absent in the first version and were added. A conventional paper structure is now used.

